# Comparative metabolomics of forest communities: Species differences in foliar chemistry are greater in the tropics

**DOI:** 10.1101/271361

**Authors:** Brian E. Sedio, John D. Parker, Sean M. McMahon, S. Joseph Wright

## Abstract

Interspecific variation in the secondary metabolites of plants constrains host specificity of insect herbivores and microbial pathogens. The intensity and specificity of these plant-pest interactions is widely believed to increase towards the Equator, leading to the prediction that secondary metabolites should differ more among co-occurring plant species in tropical communities than in temperate communities. To evaluate this prediction, we quantified metabolomic similarity for 203 tree species that represent >89% of all individuals in large forest plots in Maryland and Panama. We constructed molecular networks based on mass spectrometry of all 203 species, quantified metabolomic similarity for all pairwise combinations of species, and evaluated how pairwise metabolomic similarity varies phylogenetically. Leaf metabolomes exhibited clear phylogenetic signal for the temperate plot, with high similarity among congeneric species. In contrast, leaf metabolomes lacked phylogenetic signal for the tropical plot, with low similarity among congeners. Our results suggest that species differences in secondary chemistry comprise important axes of niche differentiation among tropical trees, especially within species-rich genera, and that the contribution of species differences in secondary chemistry to niche differences increases towards the equator in forest tree communities.

---

Recent innovations in metabolomics promise new insights into the causes of the well-known latitudinal gradient in plant diversity. Wallace (1878) and Dobzhansky (1950) proposed that biotic interactions comprise a stronger selective force than the physical environment in the tropics, and herbivore and pathogen pressure is indeed greater in the tropics than at higher latitudes (Coley and Barone 1996, Schemske et al. 2009, Lim *et al.* 2015). Ehrlich and Raven (1964) hypothesized that coevolution between herbivores and pathogens and plant defenses drives diversification of plants and their natural enemies. If latitudinal variation in selection exerted by plant enemies contributes to greater tropical plant diversity, tropical plants should be better defended and have more variable defenses than temperate plants. Several authors have tested this prediction with respect to quantitative investment in chemical defenses, such as tannins and phenolic compounds (e.g. Coley and Aide 1991), but a recent meta-analysis (Moles *et al.* 2011a) and a multi-site empirical study (Moles *et al.* 2011b) found no support for the prediction that tropical plants are better defended.

Qualitative differences in the small-molecule metabolite profiles, or metabolomes, of plants may play an important role in generating and maintaining species diversity by constraining the host ranges of plant enemies. Herbivore host ranges are narrower in the tropics than at higher latitudes (Dyer *et al.* 2007), and focused studies of tropical tree genera have found that congeneric species are often remarkably divergent in secondary chemistry (Becerra 1997, Kursar *et al.* 2009, Fine *et al.* 2013, Richards *et al.* 2015, Salazar *et al.* 2016, Sedio *et al.* 2017). Such differences in secondary metabolites may allow closely related species to carve out “niches” defined by the insects and microbes they support, and those they avoid. If the importance of biotic interactions in shaping plant communities varies over latitude, temperate and tropical forests may differ in the extent to which co-occurring species differ with respect to secondary metabolites. Few studies have considered interspecific metabolomic variation among temperate forest plants (Agrawal *et al.* 2009, Mason *et al.* 2016), and none has compared such variation in a temperate and a tropical forest at the community scale.

Many thousands of plant compounds influence their biotic interactions. The structures of most plant metabolites remain unknown (Wang et al. 2016) and any given compound is likely to be shared by few species in a community. This combination of vast chemical diversity, unknown molecular structure, and rarity of secondary metabolites has precluded the pursuit of comparative metabolomics at the large taxonomic scales necessary for the study of whole communities (Sedio 2017). However, recent innovations in mass spectrometry (MS) bioinformatics make it possible to compare the structures of thousands of unknown metabolites from diverse chemical classes in hundreds of plant species simultaneously. Here, we quantify the structural similarity of all compounds, including the many unidentified compounds (Wang *et al.* 2016). We then quantify chemical similarity for all pairwise combinations of species, incorporating shared compounds and the structural similarity of compounds unique to one species in each pair (Sedio *et al.* 2017).

We assess chemical similarity among 138 tropical and 65 temperate plant species to assess differences in chemical diversity and phylogenetic signal. We compare a tropical moist forest in Panama (9^°^ 9’ N) and a temperate deciduous forest in Maryland (38^°^ 53’ N), USA. We ask to what extent these forests differ with respect to interspecific metabolomic variation and phylogenetic signal in interspecific metabolomic variation. If the role of secondary metabolites in defining species niche differences increases toward the Equator, we expect interspecific metabolomic variation to be greater at our tropical site than our temperate site. If selection for chemical divergence increases toward the Equator, we expect chemical dissimilarity to accumulate more rapidly and phylogenetic signal in metabolomic similarity to be weaker at our tropical site than our temperate site.

## Materials and Methods

### Study Sites and Species

Barro Colorado Island (BCI), Panama (9^°^ 9’ N, 79^°^ 51’ W) supports tropical moist forest. The 2010 census of a 50-ha forest dynamics plot (FDP) recorded 301 species with individuals T 1 cm in diameter at breast height (DBH) (Condit (1998). We sampled 138 species, including the 48 most abundant species, and every species in seven of the eight most species-rich woody genera (*Eugenia* (4 species), *Inga* (17), *Miconia* (12), *Ocotea* (9), *Piper* (11), *Protium* (5) and *Psychotria* (21)). Several of these species-rich genera are paraphyletic but form monophyletic clades when subsidiary genera are merged (Erickson *et al.* 2014). Hence, these figures include *Clidemia* and *Leandra* among the *Miconia*, *Cinnamomum* and *Nectandra* among the *Ocotea* (Erickson *et al.* 2014), *Tetragastris* among the *Protium* (Fine et al. 2014), and *Carapichea* and *Palicourea* among the *Psychotria* (Nepokroeff et al. 1999). We refer to these monophyletic clades by the most species-rich generic name on BCI. The 138 species represent 89% of the stems ≥ 1 cm DBH recorded in the 2010 census.

The Smithsonian Environmental Research Center (SERC) in Edgewater, MD (38^°^ 53’ N, 76^°^ 33’ W) supports temperate deciduous forest. The 2014 census of a 16-ha FDP recorded 69 species with individuals s 1 cm DBH. We sampled all 18 introduced species recorded in the FDP and 47 native species, including all species in the three most species-rich genera [*Carya* (3 species), *Quercus* (8), and *Viburnum* (3)]. The 47 native species represent 99% of the native stems ≥ 1 cm DBH in the FDP.

### Liquid Chromatography-Tandem Mass Spectrometry (LC-MS/MS)

We collected expanding, unlignified leaves from the shaded understory for 611 randomly chosen individuals of the 203 focal species between April and August 2014. We stored samples on ice immediately and at −80 °C within five hours. Sedio *et al.* (2017, in press) describe chemical extraction and analysis methods. Briefly, 100 mg of homogenized leaf tissue was extracted twice with 700 μL 90:10 methanol:water at pH 5 for 10 min. This solvent extracts small molecules of a wide range in polarity. Mild acidity aids the extraction of alkaloids. We used ultra high-performance liquid chromatography, electrospray ionization and molecular fragmentation, and tandem mass spectrometry (MS/MS) to analyze extracts (Sedio *et al.* in press) and the Global Natural Products Social (GNPS) Molecular Networking software to cluster the MS/MS spectra into consensus spectra that represent unique molecular structures (Wang et al. 2016). We refer to consensus spectra as compounds throughout.

Molecular networks that capture the structural similarity of unknown compounds are possible because molecules with similar structures fragment into many of the same sub-structures. Thus, the similarity of mass to charge ratio (*m/z*) of the fragments of two molecules reflects their structural similarity. We quantified structural similarity for every pair of compounds as the cosine of the angle between vectors defined by the *m/z* values of their constituent fragments (Wang *et al.* 2016). Cosine values < 0.6 are unlikely to reflect meaningful levels of chemical structural similarity and were omitted from molecular networks (Watrous *et al.* 2016). Our MS data and network can be found at http://gnps.ucsd.edu/ProteoSAFe/status.jsp?task=d1f7f083fa554f2c9608f238c1ccda0e.

### Chemical Structural and Compositional Similarity (CSCS)

Sedio *et al.* (2017) developed a metric that quantifies chemical structural-compositional similarity (CSCS) over all compounds in two species. Conventional similarity indices such as Bray-Curtis incorporate shared compounds, but ignore structural similarity of unshared compounds. In contrast, CSCS incorporates the structural similarity of compounds that are unique to each species. A simple example illustrates the implications. Compounds *x* and *y* are structurally similar (cosine ≥ 0.6). Species *A* contains compound *x* but not *y*, and species *B* contains *y* but not *x*. In this example, compounds *x* and *y* contribute zero to Bray-Curtis similarity, but make a positive contribution to CSCS based on their structural similarity.

CSCS weights every pairwise combination of compounds in two species by the product of their similarity (cosine score if ≥ 0.6 or 0 otherwise) and their proportional ion intensity in each species (Sedio *et al.* 2017). To calculate proportional ion intensities, we calculated mean ion intensities for every compound over all individuals and standardized by the summed means for each species. We calculated CSCS for all 20,503 pairs of species. We also recorded the chemical similarity between each species and its nearest neighbor in chemical space by selecting the greatest CSCS value for each species. We refer to this metric as nearest-neighbor CSCS (CSCS_nn_).

### Statistical Analyses

To generate phylogenies for each forest, we pruned the ForestGEO-CTFS mega-phylogeny (Erickson *et al.* 2014) to the 126 and 34 species present in the mega-phylogeny and our BCI and SERC data, respectively (Fig. S1). This excludes species introduced to SERC. We performed phylogenetic ANOVA with the R package ‘geiger’ (Harmon et al. 2008) to determine whether CSCS_nn_ differs between forests. To determine whether CSCS differs between forests, we performed ANOVA on random draws of independent pairs of species. CSCS differed significantly between forests if 95% of 10,000 ANOVAs were significant.

To evaluate the relationship between phylogeny and metabolomic similarity, we calculated mean CSCS for all pairs of species descended from each node in the phylogeny. We refer to this metric as CSCS_mrca_, where MRCA refers to most recent common ancestor. Figure S2 illustrates this calculation. To evaluate phylogenetic signal, we regressed CSCS_mrca_ against log-transformed phylogenetic distance.

To test for differences in the chemical space occupied by two groups of species, we first used non-metric multidimensional scaling (NMDS) to reduce the molecular network to two dimensions (using the ‘MASS’ package in R, Venables and Ripley 2002). We then compared the observed difference in area occupied by the two groups with the distribution of differences generated by10,000 randomizations. Randomizations reassigned species over columns of the pairwise CSCS matrix. The chemical space occupied by two groups differed significantly if the observed difference in area was greater than 95% of randomized differences.

All analyses excluded the 18 introduced species at SERC. Appendix S1 presents results of analyses that include the introduced species.

## Results

We detected 126,746 compounds, ranging from 107.06 to 2,174.66 Daltons (Da), in foliar extracts of 185 native species from BCI and SERC. The GNPS database of natural products (Wang et al. 2016) included 130 matches with these compounds. The matches include flavonoids, piperizines, quinoline alkaloids, indole alkaloids, and terpenoids, classes of plant secondary metabolites known to include anti-herbivore defenses (Fig. 1). Networks of compounds linked by cosine scores ≥ 0.6 ranged in size from 2 to 23,029 compounds, and 95,407 compounds had cosine scores < 0.6 with every other compound (Fig. 1). In many instances, compounds unique to one or a few species comprise subnetworks of structurally similar compounds (Sedio et al. 2017). Such clusters of structurally similar compounds may represent structural precursors or alternative products from shared metabolic pathways.

**Fig. 1.**
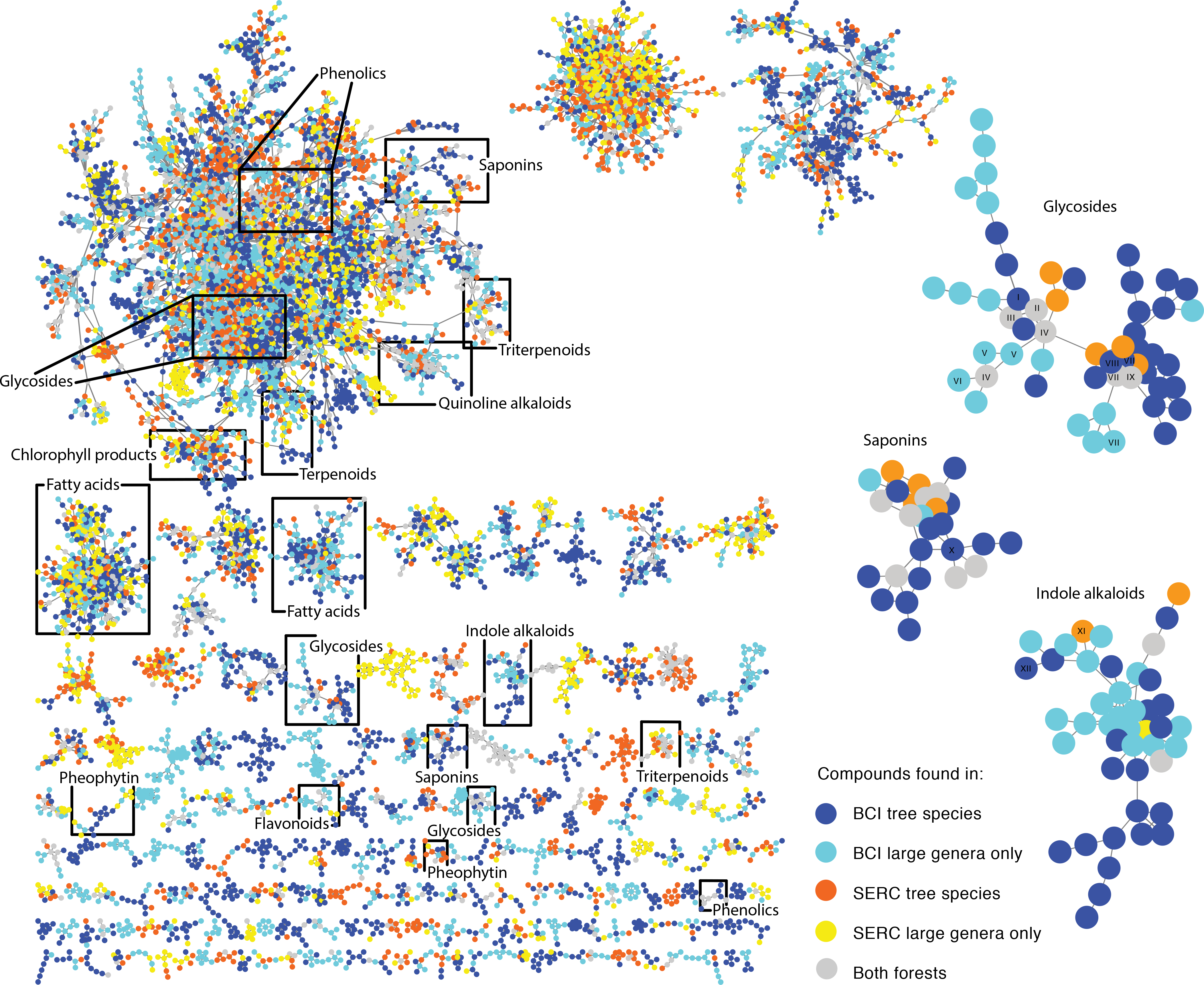
Molecular network of 36,223 compounds in leaves of tree and shrub species from SERC, Maryland and BCI, Panama. Nodes represent compounds. Links between nodes represent structural similarity between compounds indicated by cosine similarity scores ≥ 0.6. Colors represent compounds found 79 species in seven large genera at BCI (light blue), another 59 BCI species (dark blue), 14 species in three large genera at SERC (yellow) and another 33 SERC species (orange). The 130 known compounds identified chemical classes (e.g. ‘flavonoids’). We severed links with cosine scores < 0.8 to break the largest network into smaller networks for visualization. Three subnetworks are highlighted at right to illustrate compound matches to GNPS libraries. Matched compounds are I) ReSpect:PS043007 Puerarin, II) ReSpect:PM007810 3’-O-Methylluteolin 6-C-glucoside, III) ReSpect:PS086308 Orientin, IV) GNPS:Vitexin, ReSpect:PM007805 Isoorientin, VI) GNPS:Orientin, VII) GNPS:Hexanoside of (iso)orientin, VIII) GNPS:Pentoside of (iso)vitexin, IX) Massbank:PB006223 Vitexin-2”-O-rhamnoside, X) GNPS:Soyasaponin I, XI) GNPS:MLS000111555-01! Tetrahydroalstonine, XII) GNPS:Yohimbine.

The tropical, BCI species exhibited lower chemical similarity (p < 0.0001; Fig. 2a) and lower chemical similarity to their nearest neighbor in chemical space (PGLS ANOVA F_1,158_ = 45.78, p < 0.0001; Fig. 2b) than the temperate, SERC species. The largest genera made an important contribution to these site differences, with CSCS and CSCS_nn_ being much lower for the seven most species-rich BCI genera than for the three species-rich SERC genera (Fig. 2c-d).

**Fig. 2.**
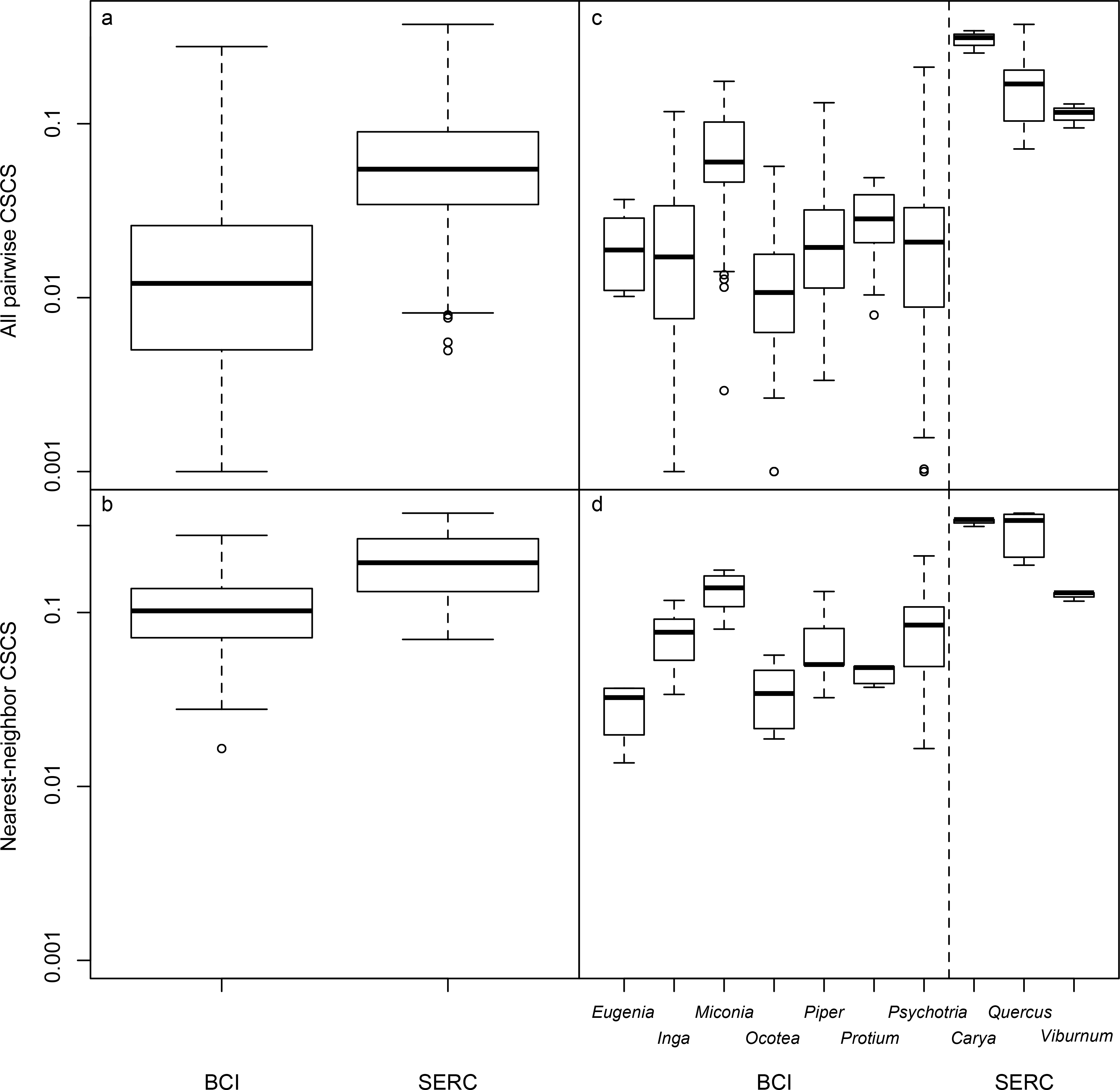
Tree and shrub species are chemically more similar at SERC and less similar at BCI. Chemical similarity for all pairwise combinations of species from BCI and from SERC (panel a) and of congeners from ten large genera (b). Chemical similarity between nearest neighbors in chemical space (CSCS_nn_) for all species from BCI and from SERC (c) and for congeners from ten large genera (d).

Among BCI species, CSCS_mrca_ was unrelated to log-transformed phylogenetic distance of most recent common ancestors (*t* = −1.28, df = 123, *p* = 0.205; Fig. 3a), indicating a strong tendency for chemical divergence among closely related species in this tropical forest. Among SERC species, CSCS_mrca_ was strongly related to phylogeny (*t* = −3.59, df = 31, *p* = 0.001; Fig. 3b), indicating that closely related species have similar metabolomes.

**Fig. 3.**
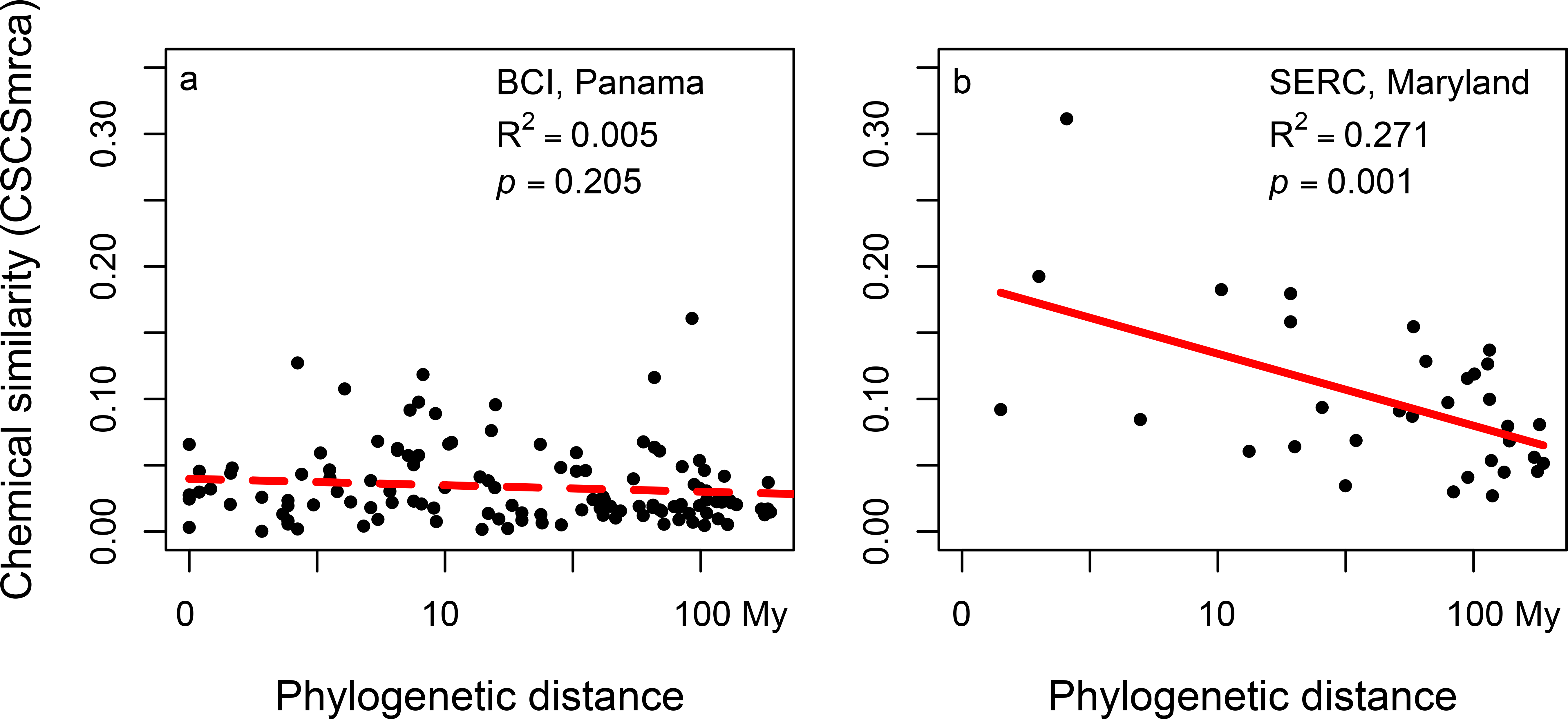
Relationships between the mean chemical similarity of species descended from each node (or most recent common ancestor, CSCS_mrca_) and log-transformed phylogenetic distance for BCI (panel a) and SERC (b). The dashed and solid red lines represent insignificant and significant linear regressions, respectively. The calculation of CSCS_mrca_ is illustrated in Fig. S2.

The NMDS ordination illustrates the chemical space represented by 138 BCI species and 47 native SERC species (Fig. 4a). Species comprising the largest BCI genera occupy a greater area in chemical space than the remaining BCI species (*p* < 0.001; Figs. 4b, 4c and 4e). In contrast, species comprising the largest SERC genera do not comprise a greater chemical space than the remaining SERC species (*p* = 0.707; Fig. 4d). Results were qualitatively similar for analyses that included the 18 introduced SERC species (Appendix S1).

**Fig. 4.**
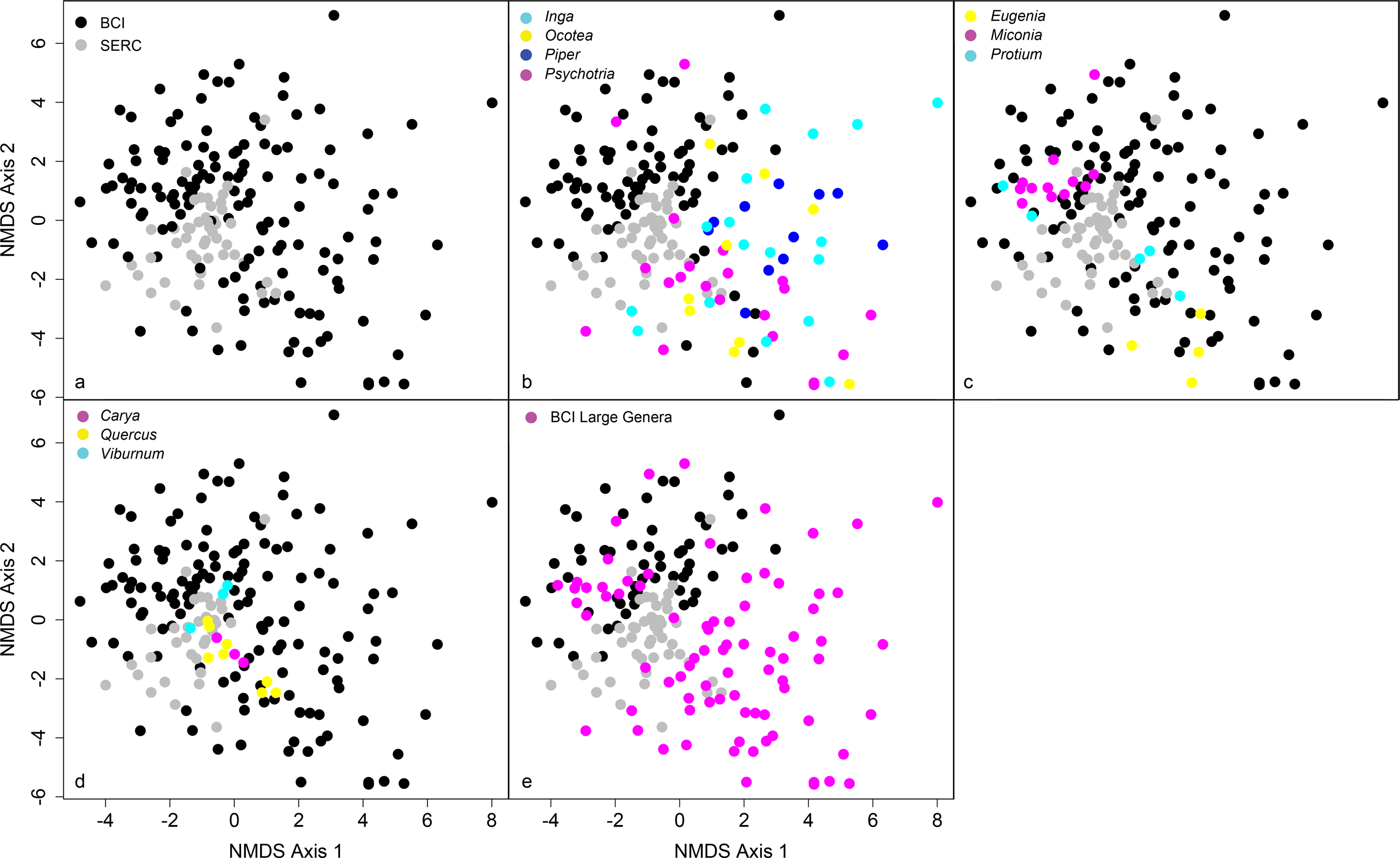
Non-metric multidimensional scaling of pairwise CSCS chemical similarity for 185 tree and shrub species. Each point represents one species, and the distances between points reflect the pairwise CSCS similarity between all pairs of species, represented in two dimensions. The 185 species include 138 species from Barro Colorado Island, Panama (black points in all panels plus colored points in panels b, c and e) and 47 native species from the Smithsonian Environmental Research Center, Maryland (gray points in all panels plus colored points in d). Colors represent seven of the largest genera at BCI (b,c), the three largest genera at SERC (d), or all seven large BCI genera (e).

## Discussion

There are fundamental chemical differences between trees from tropical, Panama and temperate, Maryland. The tropical tree species are chemically more distinctive (or dissimilar) when compared to most recent common ancestors (Figs. 3a,b), to the most chemically similar species (Figs. 2b, 4a), and over all species (Figs. 2a, 4a). Chemical similarity was also consistently lower among species-rich tropical genera than among species-rich temperate genera (Figs. 2c, 2d, 4b, 4c and 4d). These results are consistent with the hypothesis that plant-enemy interactions are more intense in the tropics, leading to rapid evolution of phytochemical diversity in tropical versus temperate trees.

The contrasting relationships between chemical similarity and phylogenetic distance for most recent common ancestors (Fig. 3) suggest contrasting selection regimes. In the BCI community, chemical similarity and phylogenetic distance are decoupled (Fig. 3a). This suggests chemical differences accrue rapidly at speciation events or with selection for divergence among closely related species. In the SERC community, chemical similarity and log-transformed phylogenetic distance are linearly related. This exponential decay of chemical similarity suggests a constant rate of chemical divergence over time. This marked contrast in phylogenetic signal suggests that selection for chemical divergence among close relatives is stronger in the tropical community and weaker in the temperate community.

The absence of phylogenetic signal in foliar metabolomic similarity among BCI tree species presents a stark contrast with leaf functional traits such as mass per area; tissue density; lamina toughness; vein toughness; cellulose, lignin, nitrogen, phosphorus and potassium content; and carbon-to-nitrogen ratio, all of which exhibit phylogenetic signal (Lebrija-Trejos *et al.* 2014). This contrast suggests that leaf chemical traits diverge more rapidly than leaf functional traits during or shortly after speciation in tropical trees and is consistent with the hypothesis that reciprocal coevolution between plants and their enemies promotes diversification, especially at low latitudes (Ehrlich and Raven 1964, Schemske *et al.* 2009).

The hypothesis that biotic interactions are more intense in the tropics and contribute to the global latitudinal diversity gradient has seen much recent controversy (Moles et al. 2011a,b). A key prediction of this hypothesis is that plants should be better defended at lower latitudes (Schemske et al. 2009). Recent evaluations of this prediction have focused on quantitative investment in defense, with mixed results (Coley and Aide 1991, Moles et al. 2011a,b). In contrast, our data suggest that qualitative chemical differences are greater among tropical species than among temperate species. Qualitative differences in chemical defenses have the potential to constrain the host ranges of herbivores and pathogens, enabling enemy-based niches, and may be especially important among members of species-rich tree genera that otherwise share similar niches (e.g. Kursar *et al.* 2009, Sedio *et al.* 2012). These qualitative differences evolved more rapidly for a tropical community than a temperate community (Fig. 3). Thus, our results suggest selection for divergence in secondary metabolites is greater in tropical than in temperate plants, even if quantitative investment is not (e.g. Moles *et al.* 2011a,b).

The extension of our conclusions beyond one tropical and one temperate forest to understand global ecological patterns will require comparative forest metabolomics of multiple sites along broad latitudinal gradients using consistent methods. Ideally, these sites would include several biogeographic regions. By enabling the study of hundreds of thousands of metabolites in hundreds of plant species, the forest metabolomic approach presented here promises to enable a more mechanistic understanding of the role that interspecific chemical variation plays in niche partitioning among co-occurring species and in lineage diversification at community, biogeographic, and macroevolutionary scales (Sedio 2017). Ultimately, integrating forest metabolomics with plant-enemy associations, recruitment dynamics, and phylogeny over geographically diverse sites will provide a critical test of the hypothesis that chemically mediated biotic interactions are a primary contributor to global patterns of plant diversity.

## Acknowledgements

We thank J.C. Rojas Echeverri, J. Trejo, J. Adams, B. Hostetler, N. Khosla, C. López, Z. Mijango Ramos, D. Plant, A. Sierra, K. Uckele, and J. I. Wright for assistance in the laboratory, P. Dorrestein, M. Meehan, R. Gittens, and C. Boya for valuable discussion. This work was supported by the Smithsonian Institution Grand Challenges Award and Scholarly Studies grant programs and a Smithsonian Tropical Research Institute Earl S. Tupper Fellowship.

